# Telomere-to-telomere genome assembly of an allotetraploid pernicious weed, *Echinochloa phyllopogon*

**DOI:** 10.1101/2023.05.23.541891

**Authors:** Mitsuhiko P. Sato, Satoshi Iwakami, Kanade Fukunishi, Kai Sugiura, Kentaro Yasuda, Sachiko Isobe, Kenta Shirasawa

## Abstract

*Echinochloa phyllopogon* is an allotetraploid pernicious weed species found in rice fields worldwide that often exhibits resistance to multiple herbicides. An accurate genome sequence is essential to comprehensively understand the genetic basis underlying the traits of this species. Here, the telomere-to-telomere genome sequence of *E. phyllopogon* was presented. Eighteen chromosome sequences spanning 1.0 Gb were constructed using the PacBio highly-fidelity long technology. Of the 18 chromosomes, 12 sequences were entirely assembled into telomere-to-telomere and gap-free contigs, whereas the remaining six sequences were constructed at the chromosomal level with only eight gaps. The sequences were assigned to the A and B genomes with total lengths of 453 and 520 Mb, respectively. Repetitive sequences occupied 42.93% of the A genome and 48.47% of the B genome, although 32,337, and 30,889 high-confidence genes were predicted in the A and B genomes, respectively. This suggested that genome extensions and gene disruptions caused by repeated sequence accumulation often occur in the B genome before polyploidization to establish a tetraploid genome. The highly accurate and comprehensive genome sequence would contribute to elucidating the population structure of this species and could be a milestone in understanding the molecular mechanisms of the pernicious traits and to developing effective weed control strategies to avoid yield loss in rice production.

## Introduction

*Echinochloa phyllopogon* (2n = 4x = 36) is a member of the Poaceae family, close to *Setaria italica*, an autogamous plant, and is a noxious weed in flooded rice worldwide. Among the *Echinochloa* genus, it is widely recognized as the species that best adapts to paddy fields (Yamasue 2001). *E. phyllopogon* was found only in watered environments, although populations have recently been discovered in paddy levees, roadsides, and other places (Yasuda et al. 2020). Herbicides have been used to manage this species owing to their huge impact on rice yields. However, repeated use of herbicides has resulted in the evolution of herbicide resistance in rice fields, posing a serious threat to agriculture.

This species often exhibits resistance to multiple herbicides, which has been attributed to the overexpression of herbicide-detoxifying enzymes such as cytochrome P450 monooxygenases (Iwakami et al. 2014, 2019; Suda et al. 2022). However, the precise molecular mechanisms underlying resistance are not yet fully understood. A highly accurate genome sequence would help elucidate these mechanisms and provide a deeper understanding of how multiple herbicide resistance occurs.

Despite the complex genome structures, including polyploidy, several species of genome sequences in *Echinochloa* were determined (Ye et al. 2020; Wu et al. 2022). Wu et al. (Wu et al. 2022) revealed the complex and reticulate evolution in the speciation of *Echinochloa* polyploids and reported the chromosome-level genome sequences (945 Mb in length) in *E. phyllopogon*, for which Continuous Long Reads (PacBio), paired-end and mate-pair reads (Illumina), and Hi-C techniques were employed. However, the assembly is shorter than the estimated genome size of 1.0 Gb (Ye et al. 2020) and included 627 gaps (ca. 50 Mb in length), in which sequences were undetermined. Gapped genome sequences that do not cover the entire genome might miss complex genome structures, genetic bases of agriculturally important traits, and evolutionarily important variations. Owing to recent advanced long read technology, telomere-to-telomere (T2T) and gap-free genome sequences have been reported in human (Nurk et al. 2022), chicken (Huang et al. 2023), fungi (Kurokochi et al. 2022; Bowyer et al. 2022), and plankton (Bliznina et al. 2021; Giguere et al. 2022). Herein, the chromosome-level assembly of the allotetraploid genome of *E. phyllopogon* was reported. The assembly included 12 T2T and gap-free sequences of the 18 chromosomes in addition to six sequences connected at the chromosome-level with only eight gaps. The genome information from this study would contribute to weed controls to avoid yield loss in rice production and to understand weed adaptation and propagation systems.

## Methods

### Plant materials

A single line of *E. phyllopogon*, R511, sampled from California, USA (Iwakami et al. 2014), was used for *de novo* genome assembly. The R511 line was crossed the S401 line (California, USA) to generate an F5 mapping population (n = 118). In addition, seven lines (oz1804, Eoz1813, Eoz1814, core109, core113, core301, and core304) from the northeastern mainland of Japan(Iwakami et al. 2014; Yasuda et al. 2020; Suda et al. 2022) were used for whole genome re-sequencing analysis.

### *De novo* whole genome assembly

To estimate the genome size of R511, genome DNA of R511 was extracted using the DNeasy Plant Mini Kit (Qiagen, Tokyo, Japan), and the sequence library was constructed with the MGIEasy PCR-Free DNA Library Prep kit (MGI Tech, Shenzhen, China) and sequenced on the DNBSEQ-G400 (MGI Tech). The genome size of line R511 was estimated using *k*-mer distribution analysis of short-read sequences (*k* = 17) with Jellyfish software (v2.3.0) (Marçais and Kingsford 2011).

For long-read sequencing, high-molecular weight genomic DNA was extracted from the leaves of R511 using NucleoBond HMW DNA (MACHEREY-NAGEL, Dueren, Germany). Genomic DNA was prepared using the SMRTbell Express Template Prep Kit (PacBio, Menlo Park, CA, USA). Long-read sequence data were obtained using a Sequel IIe system (PacBio). All HiFi reads were assembled using Hifiasm version 0.16.1 (Cheng et al. 2021) with default parameters. In parallel, a subset of HiFi reads randomly sampled from the data was assembled using Hifiasm, as described above. The two assemblies from the all datasets and the subsets were aligned with MUMmer4 (Marçais et al. 2018) to compare structures and search for contig sequences in one assembly that bridged separated sequences in another assembly.

### Chromosome-level scaffolding via genetic mapping

A genetic map of *E. phyllopogon* was established with a double-digest restriction site-associated DNA sequencing (ddRAD-Seq) technique (Peterson et al. 2012). Genomic DNA were extracted from the leaves of the F5 mapping population and its parental lines using the DNeasy Plant Mini Kit (Qiagen) and subjected to ddRAD-Seq library construction using the PstI and MspI enzymes (Shirasawa et al. 2016). ddRAD-Seq reads were obtained using HiSeq 4000 (Illumina, San Diego, CA, USA), and their low-quality bases (<10 quality value) and adaptor sequences (AGATCGGAAGAGC) were trimmed with PRINSEQ and fastx_clipper in the FASTX-Toolkit, respectively. The cleaned reads were mapped onto primary contigs constructed using all reads as a reference in Bowtie2 (Langmead and Salzberg 2012), and sequence variants were called using BCFtools (Danecek et al. 2021). High-confidence SNPs were selected using VCFtools (Danecek et al. 2011) (parameters: minDP 5, minQ 999, maf 0.2, max-maf 0.8, max-missing 0.5) and subjected to linkage analysis using Lep-Map3 (Rastas 2017). The resulting map was merged with the genome assembly using ALLMAPS (Tang et al. 2015). The contigs assembling all reads were scaffolded manually based on the downsampled assembly and genetic map, for which 100 Ns were placed between the scaffolded contigs to generate pseudomolecule sequences. Assembly quality was evaluated using BUSCO v5 with the embryophyte_odb10 data (Manni et al. 2021) and telomere sequences (TTTAGGG) were searched using telomere_finder (https://github.com/MitsuhikoP/telomere_finder). Genetic distance was calculated using the alignment-free genetic distance estimation software, Mash (Ondov et al. 2016).

### Repetitive sequence analysis and gene prediction

Repetitive sequences were detected with RepeatMasker v4.1.2 (https://www.repeatmasker.org) using repeat sequences obtained from the pseudomolecule sequence using RepeatModeler v2.0.2 (https://www.repeatmasker.org) and from a dataset registered in Repbase (Bao et al. 2015).

Potential protein-coding genes were predicted using Braker version 2.1.5 (Br na et al. 2021) with the protein sequences of *Oryza sativa* (Kawahara et al. 2013), *Zea mays* (Jiao et al. 2017), and *E. oryzicola* (Wu et al. 2022). To assign confidence to predicted genes, homologous genes against the eggNOG 5.0 (Huerta-Cepas et al. 2019) and UniProtKB databases (12 Aug 2022) (UniProt Consortium 2021) were searched using eggNOG-Mapper 2.1.8 (Huerta-Cepas et al. 2017) and DIAMOND 2.0.14 (Buchfink et al. 2021), respectively. The genes that hit against eggNOG and UniProtKB were classified as high-confidence (HC) genes; however, those that hit keywords related to transposable elements (TEs) were classified as TE. The other genes were classified as low-confidence (LC) genes. Gene clustering was performed using OrthoFinder (Emms and Kelly 2019). The mapping annotation of previously reported chromosomal sequences of *E. phyllopogon* (eo_v2) (Wu et al. 2022) was confirmed using Liftoff (Shumate and Salzberg 2020).

### Comparative genome structure analysis

Chromosome-level pseudomolecule sequences were aligned using minimap2 (v2.24) (Li 2018) and compared to closely related species using D-GENIES (Cabanettes and Klopp 2018), pafr in the R package. Synteny and collinearity of the predicted genes were detected using MCScanX (Wang et al. 2012) and visualized using SynVisio (Bandi and Gutwin 2020).

### Whole genome resequencing analysis

Genomic DNA was extracted from the leaves of the two US and seven Japanese lines using a DNeasy Plant Mini Kit (Qiagen). Genomic DNA libraries for short-read sequencing were prepared and sequenced, as described above. In addition, short-read data for 84 *E. phyllopogon* lines were obtained from a public database (Wu et al. 2022) (accession numbers CRA005291 and CRA005559), which contained two US, 55 Chinese (including nine northeast China, 39 southeast China, and seven Hainan), 25 Italian, one Korean, and one Malaysian lines. The sequence reads for 93 lines were treated as described above, and clean reads were mapped to the pseudomolecule sequence, using Bowtie2 (Langmead and Salzberg 2012); sequence variants, for example, SNPs and InDels, were called using BCFtools (Danecek et al. 2021). The variants were filtered using VCFtools (Danecek et al. 2011) (parameters: minDP 5, minQ 10, maf 0.05, max-maf 0.95, max-missing 0.8). The effects of the variants on gene function were annotated using SnpEff v4.3 (Cingolani et al. 2012). Principal component analysis (PCA) was performed for all SNPs using PLINK1.9 (Chang et al. 2015). The maximum likelihood phylogenetic relationship was inferred with synonymous SNPs using RAxML(Stamatakis 2014) with the GTRGAMMA model and 1,000 bootstraps. Population structure analysis was performed using the Admixture ver. 1.3.0 with 1,000 bootstraps.

## Results

### Genome sequence and *de novo* assembly

Based on the *k*-mer frequency analysis using short-read sequences (23.3 Gb), the genome size of *E. phyllopogon*, R511 was estimated as 1.03 Gb (Fig. 1). Subsequently, 4.5 million HiFi sequence reads (68.9 Gb; 66.9× coverage of the estimated genome size, N50 = 14.9 kb) obtained from two single-molecule real-time (SMRT) cells, were assembled into primary contigs (EPH_r1.0) and haplotigs. The assemblies consisted of 1.00 Gb of primary contigs (EPH_r1.0, including 827 sequences with an N50 length of 45 Mb, Table 1) and 50.9 Mb haplotigs (including 1,647 sequences with an N50 length of 33 kb, Table S1). The presence of a single peak in the *k*-mer distribution and the observation of short haplotigs suggest a high level of homogenetity in the genome. In the primary assembly, 38 telomere repeat sequences (TTTAGGG) were detected at the ends of 26 contigs. Of these, 12 and 14 contigs had telomere repeat sequences at both ends and at one end, respectively (Table 2). These results suggested that the 12 contigs were assembled at a gapless telomere-to-telomere (T2T) level.

**Figure 1.**
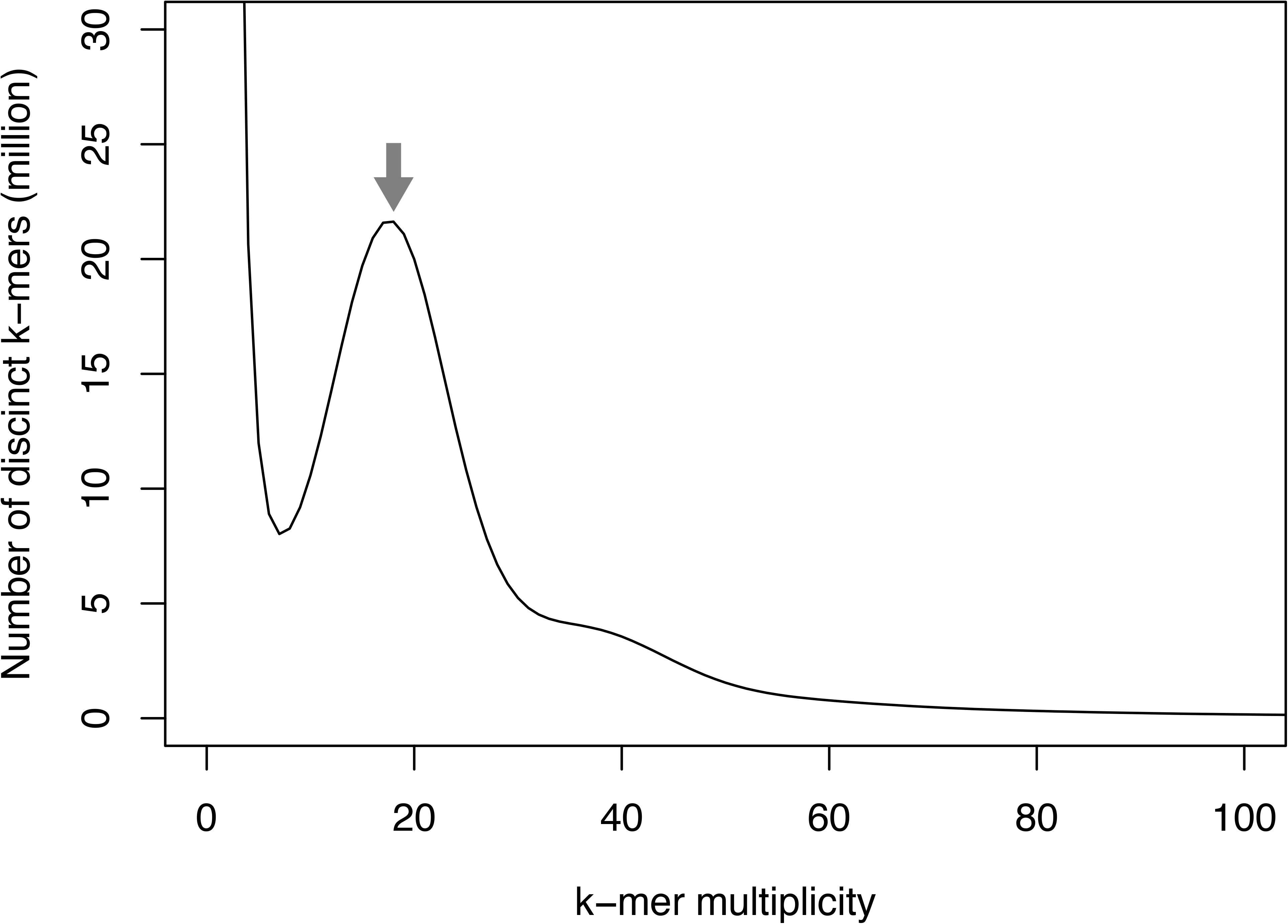
Estimation of the genome size of *E. phyllopogon*, based on *k*-mer analysis (*k* = 17) with given multiplicity values. Gray arrow indicates the peak used by estimation.

**Table 1.** Statistics of the genome assembly of *E. phyllopogon*.

|  | EPH_r1.1 | EPH_r1.0 |
| --- | --- | --- |
| Total scaffold size (bp) | 1,002,757,761 | 1,002,676,961 |
| Number of scaffolds | 19 | 827 |
| Number of chromosomes | 18 | 18 |
| Scaffold N50 length (bp) | 58,069,259 | 45,224,446 |
| Longest sequence size (bp) | 72,873,931 | 67,065,931 |
| No. of gap | 808 | - |
| gap length (bp) | 80,800 | - |
| Complete genome BUSCOs | 98.9 | 98.9 |
| Number of HC genes predicted | 64,805 | - |
| Complete HC protein BUSCOs | 95.1 | - |

**Table 2.**
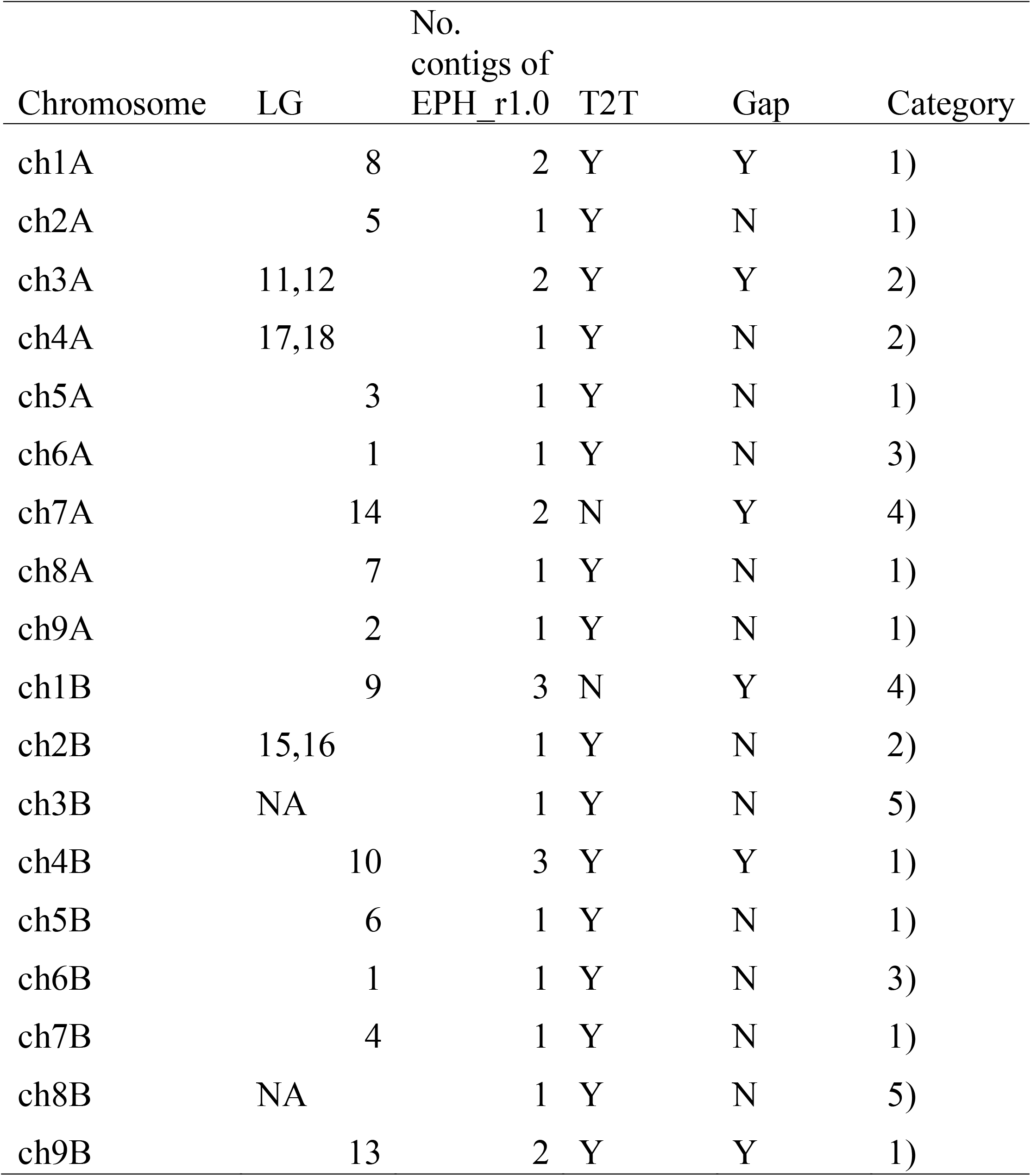
Chromosome scaffolding status using downsampling and genetic map. Chromosomes constructed via scaffolding and genetic map.

It is expected that higher coverage of HiFi reads will result in increased assembly contiguity. Whereas PacBio recommend coverages of 10 to 15-fold per haplotype for *de novo* assembly using HiFi reads (https://www.pacb.com), given very high coverage, ultra-low frequency of errors may negatively impact the contiguity (Cheng et al. 2021). A downsampling strategy was tested to determine whether low-depth reads could provide gapless T2T assemblies. Of the 68.9 Gb Hifi reads, we sampled 3 million (45.8 Gb) and established another assembly, EPH_r0.1 (Table S1). This assembly contained 580 contigs spanning 1.00 Gb with an N50 of 43 Mb. The contigs of EPH_r0.1 were aligned with those of EPH_r1.0. Six EPH_r0.1 contigs participated in scaffold 10 EPH_r1.0 contigs to generate four scaffold sequences (Fig. S1). Of these, three and one scaffold sequences contained telomere sequences at both ends and one end, respectively (Table 2).

### Chromosome-level pseudomolecule sequence construction

A genetic mapping approach was used to construct the remaining two chromosome-level sequences. First, a genetic map was constructed based on the SNPs identified using ddRAD-seq analysis (Table S2). An average of 6.0 M ddRAD-seq reads per sample were obtained, of which 95.0% were mapped onto the EPH_r1.0 assembly. After filtering by SNP-calling quality, 14,420 high-confidence SNPs were identified in 26 primary contigs; however, they were not detected in the remaining 801 contigs. A total of 12,643 SNPs were separated into 18 linkage groups and ordered to cover 2,271.6 cM in length (Table S3 and Fig. S2).

Based on genetic mapping, 18 T2T or chromosome-level contig sequences were fully constructed with the five categories (Table 2): 1) one linkage group supported one contig (in nine sequences); 2) two linkage groups covered one contig because of the absence of SNPs in the middle of the chromosomes (in three sequences); 3) one linkage group corresponded to two contigs probably because of a misjoining of the linkage map (in two sequences); 4) one linkage group corresponded to two contigs to join them into a chromosome-level contig (in two sequences); and 5) no linkage groups were constructed for contigs owing to a lack of SNPs on the entire chromosome (in two sequences). Sequences composed of multiple contigs were connected with 100 Ns. The 801 contigs without SNPs were also connected to 100 Ns to build ch00. The resulting assembly (EPH_r1.1) spanned 1,002.8 Mb in length (Table 1). The complete BUSCO score of EPH_r1.1 showed 98.9%, and the single-copy and duplicated BUSCO scores showed 9.4% and 89.5%, respectively.

To identify the A and B genomes of the tetraploid *E. phyllopogon*, 18 chromosomal sequences were compared with those of a diploid relative, *E. haploclada* (Ye et al. 2020). As expected, the nine pairs of the sequences of *E. phyllopogon* corresponded to nine chromosome sequences of *E. haploclada* (Fig. S3A). In accordance with genetic distance, sequences close to and distant from *E. haploclada* were named A and B genomes, respectively (Fig. S3B). The nomenclature and direction of the scaffold were based on nine chromosomes of foxtail millet (*Setaria italica*) (Bennetzen et al. 2012) (Fig. 2A), which are phylogenetically close and the same chromosome number to *E. phyllopogon* (Wu et al. 2022). The established pseudomolecule sequences corresponded one-to-one with previously reported chromosome sequences of *E. phyllopogon* (eo_v2) (Wu et al. 2022) (Fig. 2B).

**Figure 2.**
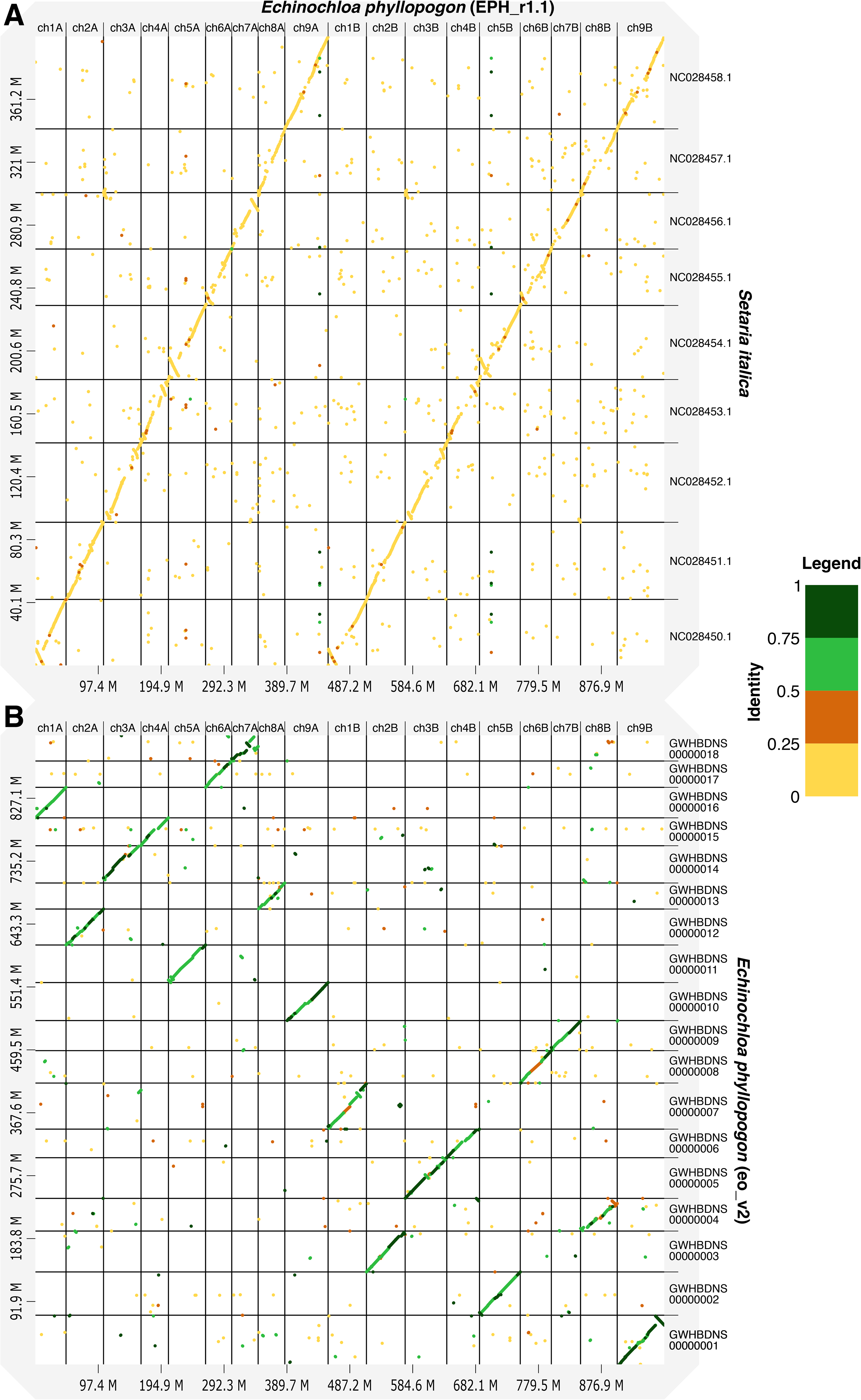
Comparative analysis of the genome sequence and structure among *E. phyllopogon* (EPH_r1.1), (A) *S. italica*, and (B) *E. phyllopogon* (eo_v2). Dots and colors indicate genome structure and sequence similarity, respectively.

### Repeat sequence analysis and gene prediction

Repeat sequences occupied 455 Mb of 1.0 Gb in EPH_r1.1 (45.40%) (Table 3). In *E. phyllopogon*, the dominant repetitive sequences were long terminal repeat (LTR) retroelements (16.46%), followed by DNA transposons (4.36%). Notably, repeat content was different between the A genome (194 Mb, 42.93%) and the B genome (252 Mb, 48.47%). The distribution of repeat sequences was similar to that of previously reported genome sequences (eo_v2) (Wu et al. 2022) except for unclassified sequences that may be unique repetitive sequences in *Echinochloa* (Table 2). This unclassified repeat in the current assembly was ∼40 Mb longer than that in eo_v2 and clustered in the middle of the chromosomes (Fig. S4) that were not represented in eo_v2.

**Table 3.**
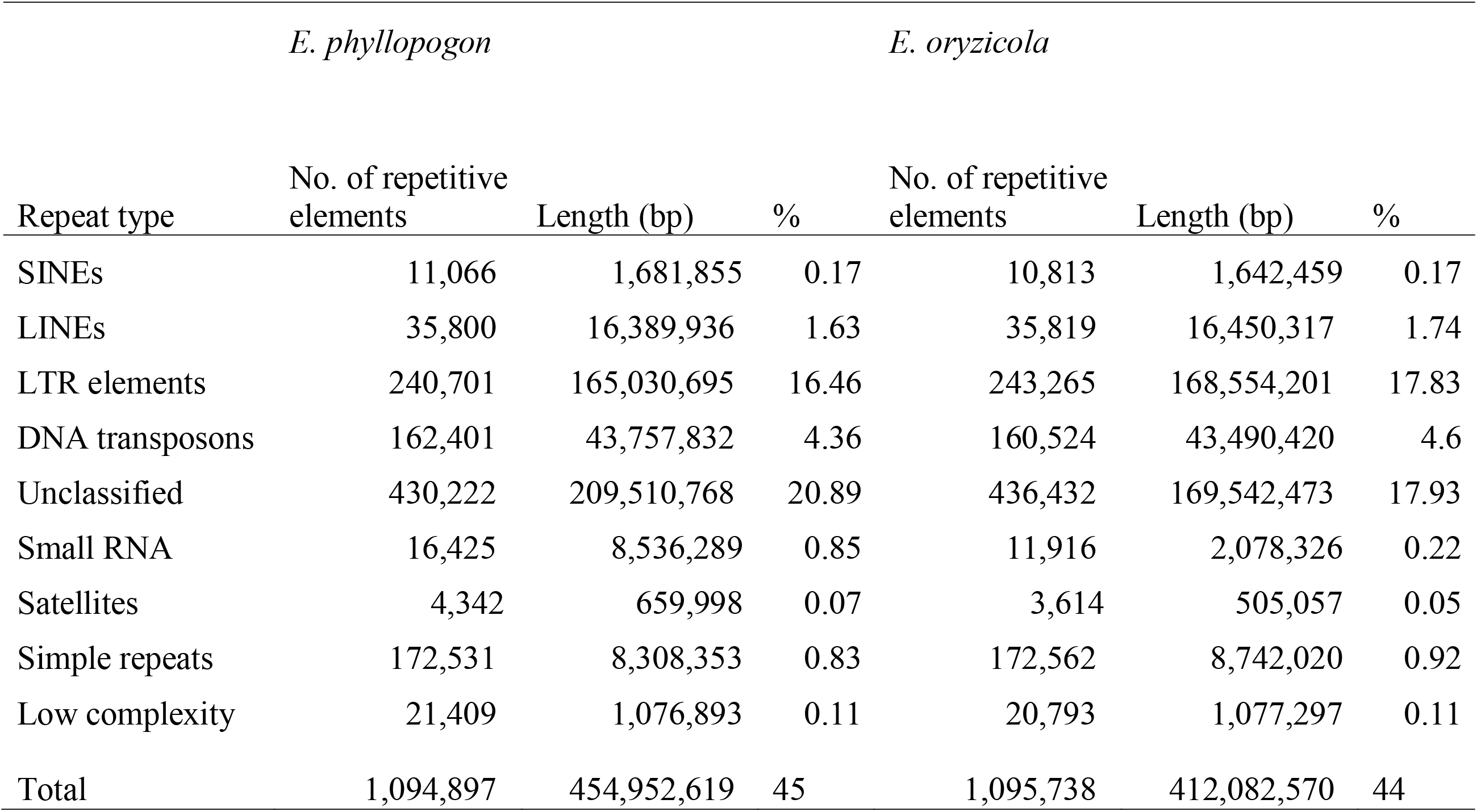
Repetitive sequences in the *E. phyllopogon* EPH_r1.1 and eo_v2 genomes.

A total of 132,212 potential protein-coding sequences were identified in the current genome assembly (Table S1), based on *ab initio* prediction and amino acid sequence homology among three Poaceae species, *O. sativa* (IRGSP-1.0) (Sakai et al. 2013), *Z. mays* (B73_v4) (Jiao et al. 2017), and *E. phyllopogon* (eo_v2) (Wu et al. 2022). In subsequent gene annotation analysis, 64,805 genes with gene descriptions were assigned as HC genes, 4,809 genes with TE-related terms were assigned as TE-related genes, and the remaining 62,598 genes were classified in LC genes (Table S4). BUSCO analysis of all and HC genes indicated that the scores for complete BUSCOs were 96.8% and 95.1%, respectively (Table S1). Of the HC genes, 97.8% (= 63,378/64,805) sequences hit the predicted genes of *E. phyllopogon* (eo_v2) (Wu et al. 2022).

A total of 32,337 HC, 2,334 TE, and 30,544 LC genes were predicted in the A genome, 30,889 HC, 2,458 TE, and 31,791 LC genes were found in the B genome (Table S1). The genes predicted in the A and B genomes of *E. phyllopogon* were clustered among the Poaceae species, *O. sativa* and *S. italica* (Fig. 3). In total, 27,120 clusters were identified. Of these, 16,144 clusters (59.5%) were shared among the four sets. A total of 1,832 clusters (343 + 1,225 + 264) were unique to *E. phyllopogon*, of which 343 and 264 clusters were unique to A genome and B genome, respectively.

**Figure 3.**
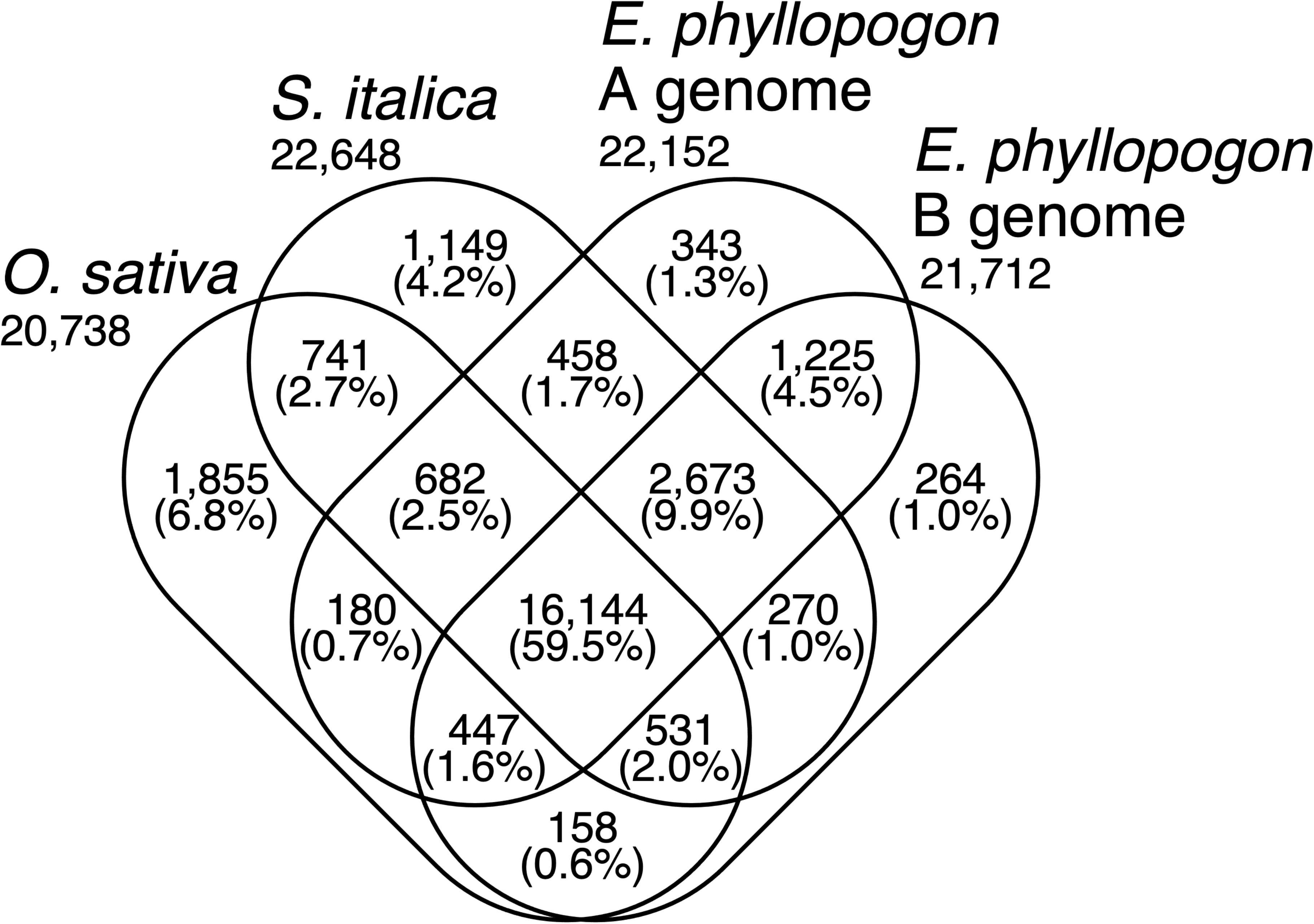
Venn diagram of the numbers of gene clusters in the A and B genomes of *E. phyllopogon* and two Poaceae species.

### Comparative analysis the genome structures of the three species

Based on sequence similarity, the structures of the A and B genomes (EPH_r1.1) and that of *S. italica* were well conserved (Fig. 2). Furthermore, we analyzed synteny based on orthologous gene orders (Fig. 4A). In sequence similarity and synteny analyses, potentially large inversions between the *E. phyllopogon* and *S. italica* genomes were observed on chromosomes 1 (inversion size of 10 Mb), 4 (6 Mb), 5 (15 Mb), 6 (10 Mb), and 7 (10 Mb) (Figs 2A and 4B). To clarify which genome had inversion events, the *E. phyllopogon* and *S. italica* were compared with those of *O. sativa* (Fig. 4). In total, 503 and 458 synteny regions were found in A and B genomes of *E. phyllopogon*, respectively, whereas 489 synteny regions were observed in *S. italica*. The synteny blocks were well conserved between *E. phyllopogon* and *S. italica*; however, some chromosomal rearrangements for *O. sativa* were found. A potential inversion on chromosome 5 was found in A and B genomes of *E. phyllopogon*, whereas inversions on chromosomes 1, 4, and 6 were found in *S. italica*. Another inversion was found on chromosome 7, which was only in A genome of *E. phyllopogon*, and not in B genome of *S. italica*.

**Figure 4.**
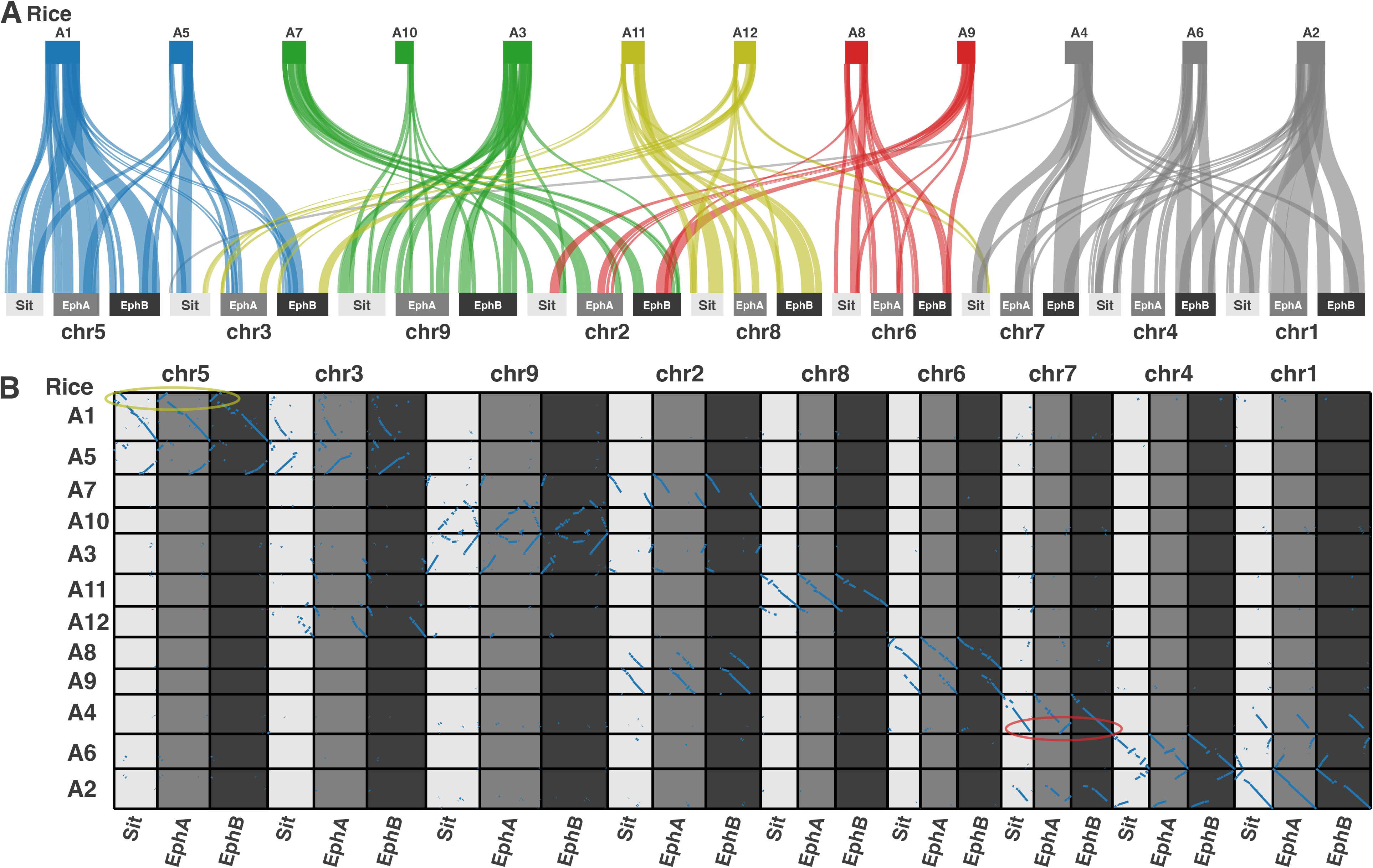
Synteny and collinearity of genes in the A and B genomes of *E. phyllopogon* (EphA and EphB) and *S. italica* (Sit) against rice chromosomes on the top. Colors of rice chromosomes indicate common ancestral genomes of Poaceae species. The yellow circle indicates *E. phyllopogon*-specific inversion and the red circle indicates A genome-specific inversion.

### Phylogenomic and population structure analysis

Phylogenomic and population structure relationships of the 93 lines were investigated using whole genome sequencing data. A mean of 131 M reads (19.71 Gb) were mapped to EPH_r1.1 as a reference, with an average map rate of 97.11% (Table S5). A total of 5,756,135 SNPs were detected across 93 lines, of which 105,949 were synonymous.

The PCA, in which the proportions of variance were 38.58% on PC1 and 9.86% on PC2, showed three clusters (Fig. 5A): A) seven Hainan lines, B) 38 Southern Chinese and one Malaysian line, and C) 25 Italian, nine Northeast Chinese, one Southern Chinese, five Japanese, four US, and one Korean line. The two Japanese lines were not included in any cluster. These clusters corresponded to those reported in a previous study (Wu et al. 2022): A) var. *hainanensis* (HN); B) Lower latitude (LL); and C) Higher Latitude (HL). Therefore, clusters A, B, and C were named HN, LL, and HL, respectively.

**Figure 5.**
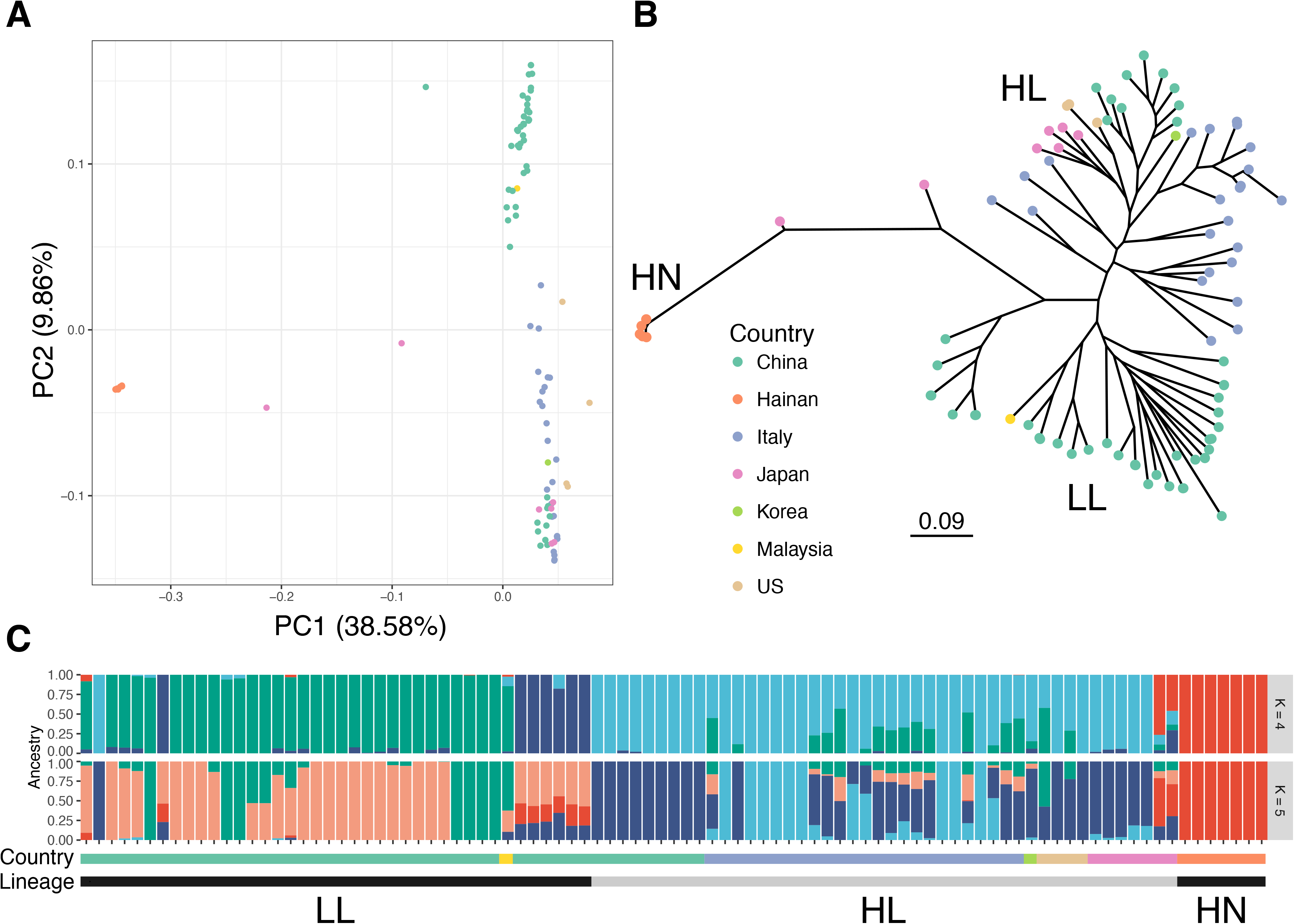
Genetic relationship for *E. phyllopogon*. (A) PCA, (B) phylogenetic tree using the maximum likelihood method, and (C) the genetic structure of K = 4 and 5. Dot colors in (A) and (B) were consistent with the horizontal bar colors in (C).

The phylogenetic tree had three main lineages–HN, LL, and HL–corresponding to the three PCA clusters (Fig. 5B). The HN group included seven Hainan lines, together with two Japanese lines that were not included in any cluster in the PCA. The HL group consisted of nine Northeast Chinese, five Japanese, and one Korean, 25 Italian, and four US lines, whereas the LL group consisted of 39 lines from 38 Southern Chinese and one Malaysian line. The US line R511, for which we determined the genome sequence in this study, belongs to HL, whereas eo_v2 belongs to LL. The population genetic structure examined using ADMIXTURE revealed that the optimal number of clusters was five (Fig. S5A). This result agrees with the PCA classifications and the phylogenetic tree. It was suggested that the two Japanese lines close to HN were potential hybrids between the lines from Hainan, Japan, and Southern China (Figs. 5C and S5).

## Discussion

Here, we present the genome assembly of the tetraploid pernicious weed, *E. phyllopogon* at the chromosome level (Tables 1 and 2). High coverage and high-quality HiFi reads could contribute to complete *de novo* assembly (Table 1). Telomeric repeat analysis revealed that 12 of the 18 pseudomolecule sequences were constructed as T2T and gap-free contigs, whereas the remaining six were constructed as chromosome-level contigs with eight gaps (Table 2). Recently, T2T genome assemblies have been reported in humans (Nurk et al. 2022), chicken (Huang et al. 2023), fungi (Kurokochi et al. 2022; Bowyer et al. 2022), and plankton (Bliznina et al. 2021; Giguere et al. 2022). In plants, gapless T2T genome assemblies have been reported in some chromosomes of maize (Liu et al. 2020), *Arabidopsis thaliana* (Wang et al. 2022), and a complete set of chromosomes of watermelon (*Citrullus lanatus*) (Deng et al. 2022). Here, we reported the chromosomal sequences of *E. phyllopogon* at the T2T level (Tables 1 and 2), even though *E. phyllopogon* has a larger and more complex tetraploid genome than those reported thus far.

The T2T genome enhances our understanding of the evolutionary processes underlying intricate genome structures, including repeat sequences, which have been historically underestimated. The 18 pseudomolecule sequences were grouped into A and B genomes in accordance with their genetic distances from a diploid relative, *E. haploclada* (Fig. S3). Comparative analysis of A and B genomes indicated that B genome (520 Mb) was longer than A genome (453 Mb), whereas the number of HC genes in B genome (30,889) was smaller than that in A genome (32,337) (Table S1). This difference was observed across all chromosomes (Fig. S4). Events of genome extension and gene disruption caused by repeat sequence accumulation (Table 3, 42.93% in A genome but 48.47% in B genome) may often occur in B genome before polyploidization to establish the tetraploid genome of *E. phyllopogon*.

The structure of the genome assembly obtained in this study was consistent with that reported in a previous study (eo_v2) (Figs. 2 and S4). The genome assembly in this study (1.0 Gb) was longer than the eo_v2 assembly (945 Mb), which was close to the estimated size of the genome of *E. phyllopogon* (1.0 Gb) (Fig. 1). The difference in length was derived from the complete assemblies of the repetitive sequences in the middle of the chromosomes, which were absent in the previously reported sequence eo_v2 (Fig. S4). Additionally, we identified structural variations between the sequences from this study and those from a previous study (Figs. 2B and S4). As the two lines belong to different clades (Fig. 5B), there may be structural polymorphisms in *E. phyllopogon*. Moreover, in a comparative analysis of genome structures among Poaceae species (Fig. 4), chromosome structure variations were uniquely found in *E. phyllopogon* and only A genome.

Phylogenomic analyses revealed at least three groups (HN, HL, and LL) of *E. phyllopogon* (Fig. 5). In HL, Northeast China, Japan, the US, and Italian lines were included, and the Italian lines were located at the base of the lineage (Fig. 5B). This suggests that the ancestor of the HL lineage first invaded Italy and subsequently expanded into Northeast China, Japan and the US, rather than parapatric differentiation within China. In the HN group, which has been reported as a new variety (Wu et al. 2022), the two Japanese lines may be hybrids of Hainan, Japan, and Southern China. The two Japanese lines possessed morphological characteristics, such as plagiotropic (or prostrate) tillers and small seed size in paddy field (Yasuda et al. 2020), which were distinguishable from those of other Japanese lines belonging to the HL group. Although it is unclear how the potential hybrids were generated and inhabited Japan, weed control to prevent migration and expansion of the novel pernicious weeds is required inside and outside paddy fields.

The highly accurate genome information of *E. phyllopogon* obtained in this study provides insights into the molecular mechanisms underlying multiple resistance to herbicides to avoid serious crop yield loss. Furthermore, *E. phyllopogon* genomics substantially contributes to the understanding of weed adaptation to farms and the pathways of worldwide weed invasion.

## Data availability

Raw sequencing reads and assemblies were deposited in the DNA Data Bank of Japan (DDBJ) under the accession number PRJDB14855. Genomic information generated in this study is available from Plant GARDEN (https://plantgarden.jp/).

## Legends

Supplementary figure 1. Dot plot comparing the alignment between EPH_r1.0, which includes all reads, and EPH_r0.1, which has been downsampled. Red lines indicate scaffolded contigs in EPH_r0.1.

Supplementary figure 2. Relationship between physical and genetic distance on the contig of EPH_r1.0. (A) Alignments and correlation between physical and genetic distance on six scaffolded contigs. (B) The marker position of contigs (EPH_r1.0).

Supplementary figure 3. Comparative analysis among *E. phyllopogon* (EPH_r1.1) and *E. hploclada*. (A) Comparison of the genome sequence and structure. Dots and colors indicate genome structure and sequence similarity, respectively. (B) Genetic distance of homologous chromosomes. *S. italica* (Sit) is outgroup. The blue letters of chromosome of *E. phyllopogon* indicate A genome and green indicate B genome.

Supplementary figure 4. Alignment region and repeat occupancy between *E. phyllopogon* EPH_r1.1 and eo_v2.

Supplementary figure 5. Population genetic structure analysis of *E. phyllopogon* worldwide. (A) Cross-valid error (B) K = 3–8.

Supplementary table 1. Detailed statistics of the genome assembly and gene prediction of *E. phyllopogon*.

Supplementary table 2. The number of reads and mapping rate of ddRAD-Seq.

Supplementary table 3. Linkage groups constructed by genetic mapping.

Supplementary table 4. Annotation list of high-confident genes.

Supplementary table 5. The number of reads and mapping rate of resequence worldwide.

## Acknowledgements

We thank Y. Kishida, S. Nakayama and A. Watanabe (Kazusa DNA Research Institute) for technical assistance. This work was supported by JSPS KAKENHI grant number 19H02955, 22H02347, 22H05172, and 22H05181.

